# The soil microbiome may offer solutions to ginger cultivation

**DOI:** 10.1101/2022.05.03.490558

**Authors:** Chih-Wei Wang, Jing-Wen Michelle Wong, Shu-Shuo Yeh, Yunli Eric Hsieh, Ching-Hung Tseng, Shan-Hua Yang, Sen-Lin Tang

## Abstract

The Taitung region is one of Taiwan’s main places for ginger agriculture. Due to issues with disease and nutrient, farmers cannot use continuous cropping techniques on ginger, meaning that the ginger industry is constantly searching for new lands. Continuous cropping increases the risk of infection by *Pythium myriotylum* and *Ralstonia solanacearum*, which cause soft rot disease and bacterial wilt, respectively. In addition, fertilizer additives cannot recover the soil when using continuous cropping on ginger, even when there is no decrease in trace elements observed in the soil. Although there may be other reasons for the reduction in production, such as soil microbes, we know little about the soil microbiome associated with ginger cultivation. Hence, in this study, we used the bacterial 16S V3–V4 hypervariable region of the 16S ribosomal RNA region to investigate microbe compositions in ginger soil to identify the difference between ginger soil with and without disease. Later, to investigate the influence of the well-known biocontrol agent-*B. velezensis* and fungicide Etridiazole on soil microbes and ginger productivity, we designed an experiment that collected the soil samples according to the different ginger cultivation periods to examine the microbial community dynamics in the rhizome and bulk soil. We demonstrated that *B. velezensis* is beneficial to ginger reproduction and suggest that it may influence the plant by adjusting its soil microbial composition. Etridiazole, on the other hand, may have some side effects on the ginger or beneficial bacteria in the soils, inhibiting ginger reproduction.

## Introduction

Ginger (*Zingiber officinale*) is an herbaceous perennial with underground rhizomes that are widely used as a fresh vegetable, spice, and herbal medicine. Its benefits include stimulating appetite, improving gastrointestinal motility, promoting sweating, promoting a healthy stomach, reducing cold symptoms, refreshing the mind, and reducing unpleasant seafood odor to enhance the flavor and aroma of seafood cuisine. According to statistics from the Taiwan Agriculture and Food Agency, Council of Agriculture, Executive Yuan (AFA), Taitung County has more than 200 hectares for ginger farming, making it the second-largest ginger producing area in Taiwan.

The major diseases that threaten ginger production include the soft rot disease caused by *Pythium myriotylum* and ginger bacterial wilt from the bacterial pathogen *Ralstonia solanacearum*. Soft rot disease tends to be prevalent during the summer or autumn months as *P. myriotylum* prefers warmer weather and wet soil conditions. *P. myriotylum* produces numerous mobile spores, moves through flowing water from rainfall or irrigation, and spreads rapidly from infected regions to the entire ginger plant. Based on Stirling, G.R.’s (2009) (1) report, it can take as little as 2 months for soft rot disease to infect the entire plant and completely destroy a harvest. The transmission of bacterial wilt is similar to soft rot disease, but takes longer to progress and may occur at any time during the growth period, causing huge losses in yield.

The first step to preventing the diseases is to get healthy, pathogen-free seeds. Ginger reproduces asexually using the rhizomes harvested from the previous farming period as the mother rhizomes; these are cut into pieces to make seed rhizomes. Experienced farmers will choose the mother rhizomes collected in the previous year from disease-free land. In addition, most farmers need to find a new land that has never had ginger planted in it to reduce the risk of disease. However, lands that are suitable for ginger are becoming more and more difficult to find and farmers will sometimes farm illegally in forests or developed land, affecting soil and water conservation and homeland security. Moreover, once the disease becomes widespread, the consequent reduction in ginger yield can cause serious economic losses. This is a major challenge to overcome in ginger agriculture.

This problem of ginger crop reduction cannot be improved with fertilizer (Personal communications with ginger farmers). Soil analysis showed no significant changes in the concentration of microelements (C. W. Wang, unpublished data), meaning that there are other factors that lead to reductions in yield. It is currently speculated that long-term application of chemical pesticides and fertilizers change the soil microbiome, preventing the continuous cropping of ginger and promoting disease occurrence, but the exact cause of ginger cultivation issue is still unknown. Previous studies on ginger explored ways to promote ginger growth by investigating the plant’s physiological properties and revealing the biosynthetic pathway of bioactive compounds and their benefits to human health. However, little is understood about the obstacles to ginger cropping or the plant’s soil microbiome.

Recently, fast, cost-efficient and convenient next generation sequencing (NGS) technology has made it easier to explore genome sequences and reveal gene expressions responsible for the biosynthesis of bioactive compounds in ginger. Studies on the ginger rhizome based on *de novo* transcriptome analysis, genome sequencing, and metabolomic analysis have provided molecular information on bioactive compounds stored in ginger rhizomes such as gingerol, volatile oil, and diarylheptanoids (2) and identified the 12 enzymatic gene families that are involved in the biosynthesis of gingerol (3).Furthermore, based on studies in India, comparative transcriptome analysis in *Zingiber officinale* Rosc. and *Curcuma amada* Roxb. have yielded information about genes related to the resistance mechanism against bacterial wilt infection (4) while the effect of different agro-climatic conditions on ginger secondary metabolism expression have also been identified (5).

In addition, plant growth-promoting rhizobacteria (PGPR) are widely used in agriculture, such as Gram-positive *Bacillus* species, because they are associated with many crops and form an endospore during hot and dry conditions. For example, *B. velezensis* is a member of the genus *Bacillus* and acts as a powerful biocontrol agent in agriculture. It has been used as a biocontrol agent against *Ralstonia solanacearum*, which causes tomato and banana *Fusarium* wilt disease (6). Although *B. velezensis* has been widely used to promote plant growth, whether *B. velezensis* improves ginger growth and whether it changes the root-associated microbiome are still open questions.

Although there have been many studies on the microbial compositions of crops, the difference between a healthy and diseased ginger soil microbiome remains unclear. The dynamics in the soil microbiome of ginger after adding chemicals and biocontrol agents is also unknown. Therefore, there are two parts to this study. First, to understand the microbial community dynamics in the soil close to the ginger roots (rhizosphere-detritusphere habitats) and the bulk soil, we compared the microbial composition in soils of healthy ginger and diseased ginger. In the second part, to investigate the influence of PGPR and fungicide on the soil microbes and productivity of ginger, we designed an experiment that collected soil samples according to the different ginger cultivation periods to examine the microbial community dynamics in the soil close to the ginger roots and the nearby soil after adding *B. velezensis* or the fungicide Etridiazole.

## Materials and Methods

### Collecting soil samples from diseased ginger

The soils of 32 ginger plants showing yellowing and wilting symptoms were collected from 12 different ginger fields in Taitung County from July 3 to October 21, 2019. For each diseased ginger plant, the bulk soil and rhizome soil were collected. All of the soil samples were stored in a -80°C freezer until DNA extraction.

### Treatment experiment of *B. velezensis* or fungicide etridiazole

The experimental field (GPS: 22.940611°N; 121.123861°E) is located in Luye Township, Taitung County, Taiwan. Ginger plants were planted on March 29, 2019, using four treatments: BV138 200x dilution (BL group), BV138 25x dilution (BH group), Terrazole® (containing 25% etridiazole) 1500x dilution (Etr group), and a control irrigated with water. Each treatment plot was 3 meters long and 0.3 meters wide, with two rows planted with 40 ginger seed rhizomes. Each treatment was conducted with four replicates and arranged by Randomized Complete Block Design (RCBD). The BV138 microbial reagent used in the experiment was isolated from the soil in Taitung and identified as *Bacillus velezensis*. The BV138 was manufactured by Yuan-Mei Biotech (Taichung, Taiwan) as powder containing 5×10^9^ CFU/g of *B. velezensis* cells. For each BV138 treatment, 12.5 L of diluted reagent was irrigated.

The reagents were added on April 3, May 2, June 5, June 18, July 3, July 18, July 31, August 16, August 21, and September 11, 2019. When collecting soil samples, 40 ginger plants in each treatment were randomly selected, uprooted, and shaken vigorously to remove soil attached to the rhizomes and roots. Soil was collected from the planting site down to 15 cm deep and labelled as bulk soil. The soil tightly attached to the rhizomes and roots was brushed off with a sterilized paintbrush and collected as the rhizome soil. All of the soil samples were stored in a -80°C freezer until DNA extraction. The soil samples were collected on April 26, June 3, July 1, August 26, and November 13, 2019, and January 6, 2020. The soil samples collected from six randomly picked ginger rhizomes in the same field prior to irrigation on April 3, 2019 were defined as “Day 0” of the experiment (12 samples). The experimental field was managed regularly by a farmer with over 30 years of experience cultivating ginger. Herbicide was applied on April 1, 2019. Fungicide and pesticide were applied on May 14, June 30, July 20, August 23, September 15, and September 28, 2019. Fertilizer was applied on June 16, July 26, September 15, and October 18, 2019.

## DNA extraction, marker gene amplification, barcoding, and sequencing

DNA extraction was performed using the DNeasy PowerSoil Kit (QIAGEN, MD, USA) according to the manufacturer’s protocol. For the bacterial composition survey, the V3–V4 hypervariable region of the 16S ribosomal RNA (rRNA) genes was amplified using polymerase chain reaction (PCR) with the primers 341F (5’-CCTACGGGNGGCWGCAG-3’) and 805R (5’-GACTACHVGGGTATCTAATCC-3’). Subsequently, DNA tagging PCR (five cycles) was used to tag each of the PCR products (every six samples were tagged individually and mixed). The PCR products were run in 2% agarose gel (SeaKem LE Agarose, Lonza, ME, USA), purified with a MinElute Gel Extraction Kit (Qiagen, Hilden, Germany), and quantified using a QuantiFluor dsDNA System (Promega Corporation, Madison, WI, USA) on a Qubit 2.0 Fluorometer (Invitrogen, Grand Island, NY). The paired-end library was constructed with a Celero DNA-Seq System (1-96) (Nugen, San Carlos, CA, USA); all procedures were in accordance with the manufacturers’ instructions. The library concentration and quality were assessed on a Bioanalyzer 2100 (Agilent Technologies, Santa Clara, CA, USA) using a DNA 1000 lab chip (Agilent Technologies). 16S amplicon libraries were sequenced 2 × 301 + 16 bp (dual index) using a Miseq Reagent kit v3 (600 cycles) on an Illumina MiSeq system. PCR, barcoding, and sequencing experiments were performed by Tri-I Biotech (New Taipei City, Taiwan). All of the bacterial community sequences were deposited into GenBank (SRA accession: PRJNA826673).

## Bioinformatic analyses and statistics

The 16S rRNA gene amplicon sequences were processed using the Quantitative Insights Into Microbial Ecology 2 (QIIME 2) pipeline (version 2019.10) (7). The raw reads were flipped into the same orientation by cutadapt (version 1.15) (8), truncated at 235 bp at both ends, and denoised using the DADA2 plugin of QIIME2 (9). The amplicon sequence variants (ASVs) were obtained via the denoising process with quality filtering and chimera removal. ASV taxonomy was assigned using the classifier-consensus-vsearch plugin (10) against SILVA NR132 99% 16S rRNA gene sequences (11, 12). The ASVs of chloroplast and mitochondria were removed, and then the dataset was rarefied at the minimal read counts among samples (3250 reads).

The soil bacterial community analyses were conducted and visualized using the MARco (13), vegan(14), and pheatmap (15) packages in R software (16). A Kruskal-Wallis test and Dunn’s post-hoc test were used for all statistical analyses of group comparisons with a significance level of α=0.05, and the p values were adjusted with a false discovery rate (FDR). Alpha diversity indices were estimated by richness, Shannon’s index, Simpson’s index and Chao1 index. Beta diversity of microbial communities was measured by Bray-Curtis dissimilarity using a principal coordinates analysis (PCoA), and heterogeneity was tested using ADONIS and ANOSIM tests.

## Results

### Comparison of diseased and healthy soils

By comparing the microbial composition of different diseased and healthy soil samples, we observed that the microbial composition was distinct among healthy soil, soft rot disease soil, and ginger bacterial wilt soil samples (Figure 1, ANOSIM R=0.306, p=0.001). When focusing on the rhizome part, the microbial compositions of three different soil samples were significantly different (ANOSIM R=0.357, p=0.001), while the bulk of the microbial composition had only minor differences (ANOSIM R=0.261, p=0.001) compared to that of the rhizome part. In healthy soil, remarkable similarities were found between the bulk and rhizome soil microbial compositions, but in diseased soil, the microbiome compositions of the bulk and rhizome parts were different (Figure 1), showing that the microbial composition of bulk and rhizome part of ginger changed after being infected.

**Figure 1.**
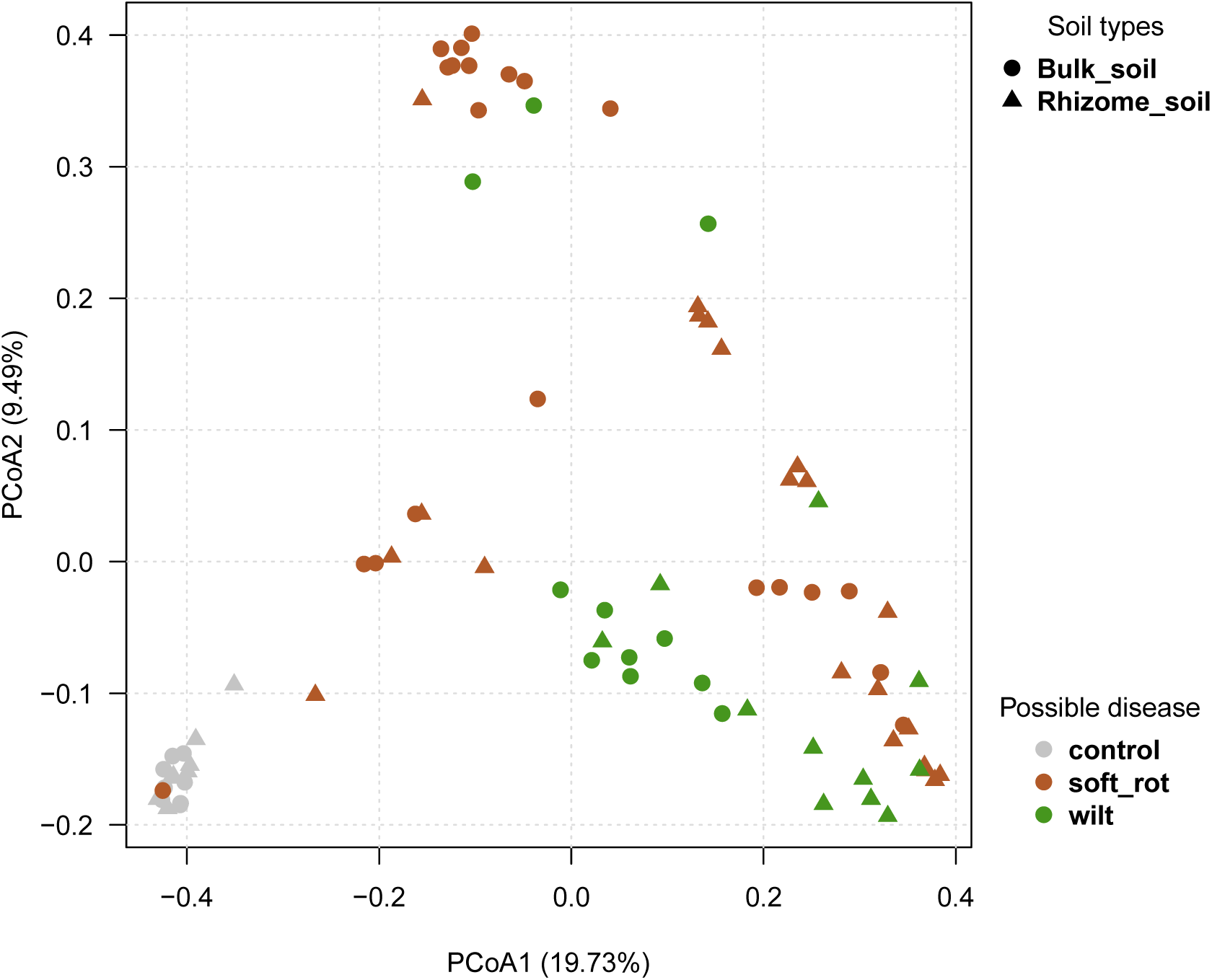
Principal Coordinates Analysis (PCoA) of bacterial communities from bulk and rhizome parts of healthy soil and two diseased soils. The forms indicate bulk soil and the colors indicate healthy and diseased soils.

The rhizome part had lower bacterial diversity than the bulk part in the healthy soil, but the difference was not significant (Figure S1). The results suggest that bacterial diversity decreased after the ginger was infected, and the infected area was mainly limited to the soil that the ginger root came into contact with. Based on figures 1 and S1, the microbial composition of the ginger rhizome part changed and its bacterial diversity was lower than in the ginger bulk part.

The 30 most abundant bacterial genera (Top 30 genera) in the rhizome parts of the healthy and diseased soils showed that the two soil types had remarkably different microbial compositions (Figure 2). In the diseased soil samples, all the ginger bacterial wilt soils were dominated by the genus *Ralstonia*, and most of the soft root disease soil was also enriched by *Ralstonia*. The relative abundance of *Ralstonia* was low in healthy soil. *Arcobacter, Dysgonomonas, Pectobacterium*, and *Myroides* were also found in diseased soil but not in healthy soil. In addition, *Flavobacterium* was the dominant bacterial genus in diseased soil, but not in healthy soil. Additionally, relative abundances of *Bacillus, Sphingomonas, Acidibacteri, Nitrosomonadaceae, Pedomicrobium, Thermoanaerobaculaceae, Nocardioides, Actinoplanes, Dongia, Terrimonas, Bryobacter, Phycisphaeraceae*, and *Steroidobacter* were greater in healthy soil than diseased soil (Figure 2).

**Figure 2.**
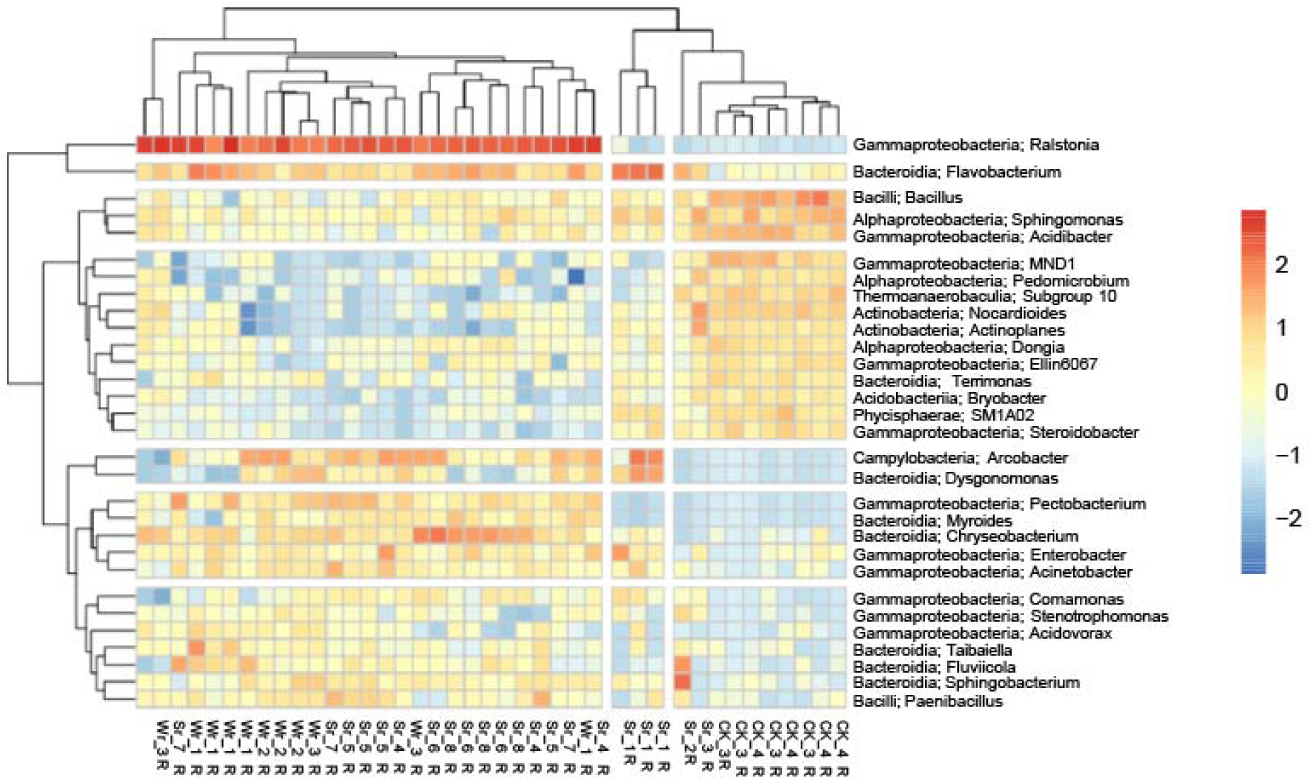
The heatmap of two-ways clustering of the top 30 genera in the rhizome part of healthy and diseased soils. The colors indicate relative abundance: warmer colors (yellow to red) indicate higher abundances and cooler colors (blues) indicate lower abundances. CK indicates control soil samples, Sr is soil with soft rot disease, and Wr is soil with wilt disease.

### Treatment experiment with *B. velezensis* or fungicide etridiazole

In the dosing test, comparing all the results of each treatment, the microbial composition changed over time (Figure 3). Including the control group, each group changed over time. At the final time point, the Etr group was distinct from the other treatment groups (Figure 3). Based on the results, time was the major factor influencing the microbial composition, but the effects varied in different treatments. Since the previous diversity index results showed that the microbial composition of the rhizome part changed greatly, the change in the microbial composition of the rhizome part in each treatment group was analyzed. As a result, the microbial composition of the rhizome part of each treatment group changed over time (Figure S2, ANOSIM: in control, R=0.637, p=0.001; in BL, R=0.795, p=0.001; in BH, R=0.728, p=0.001; in Etr, R=0.687, p=0.001).

**Figure 3.**
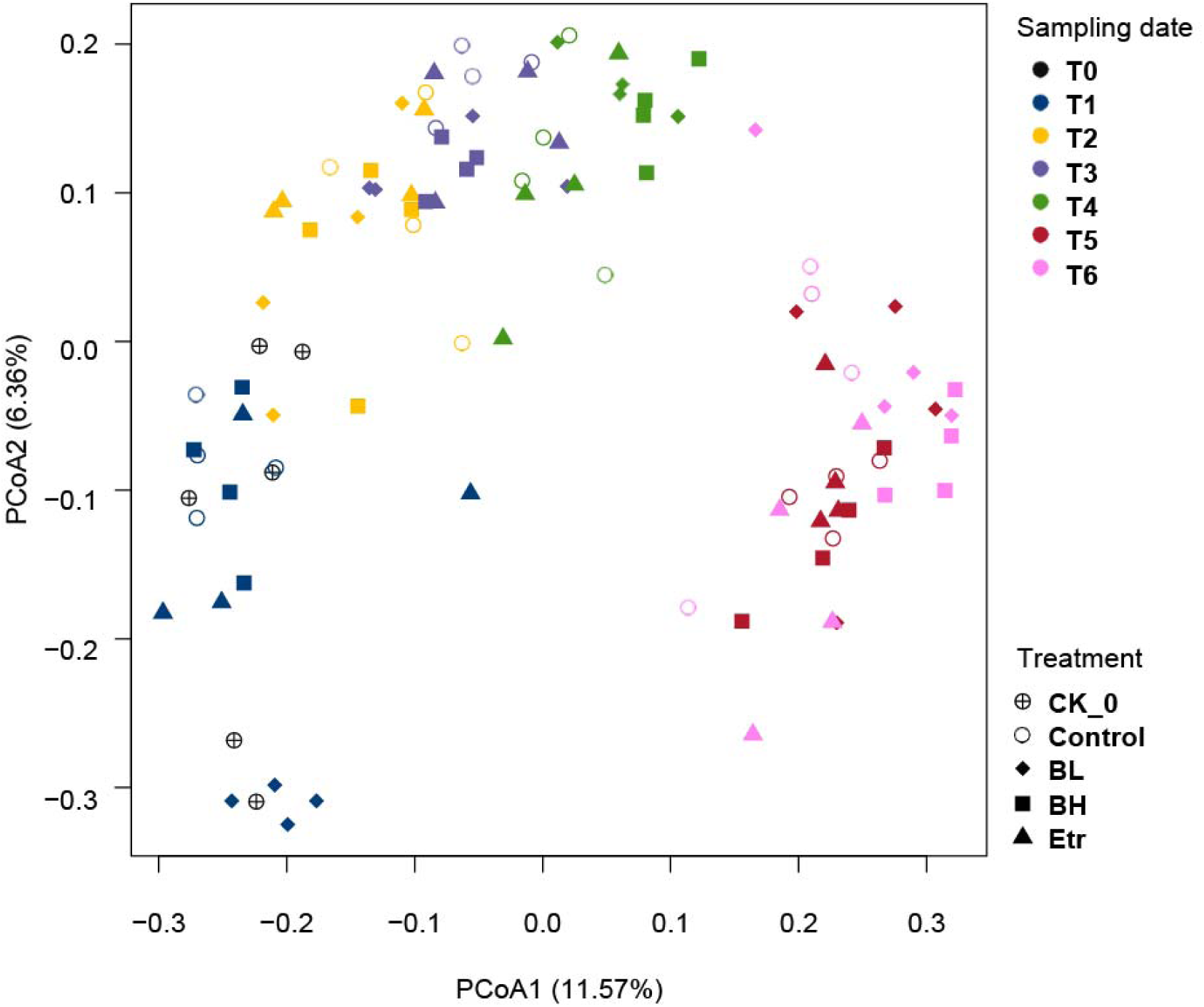
PCoA of bacterial communities from rhizome soils with different treatments along sampling times. The shapes and colors indicate different treatments and sampling times, respectively. CK0 indicates the soil before the experiment began. Control indicates ginger soil without any treatment, BL is low amounts of *B. velezensis*, BH is high amounts of *B. velezensis*, and Etr is the Etridiazole treatment.

Based on the 30 most abundant ASVs in the rhizome part of each treatment group at every time point, we observed that the dominant microbial composition changed continually (Figure S3). At different time points, we observed a remarkable similarity between BH and BL microbial compositions, while the Etr group bacterial composition was similar to that of the control group (CK). This situation continued until the final time point (Figure 4), when the BH and BL groups were clustered and the Etr group was clustered with the control group.

**Figure 4.**
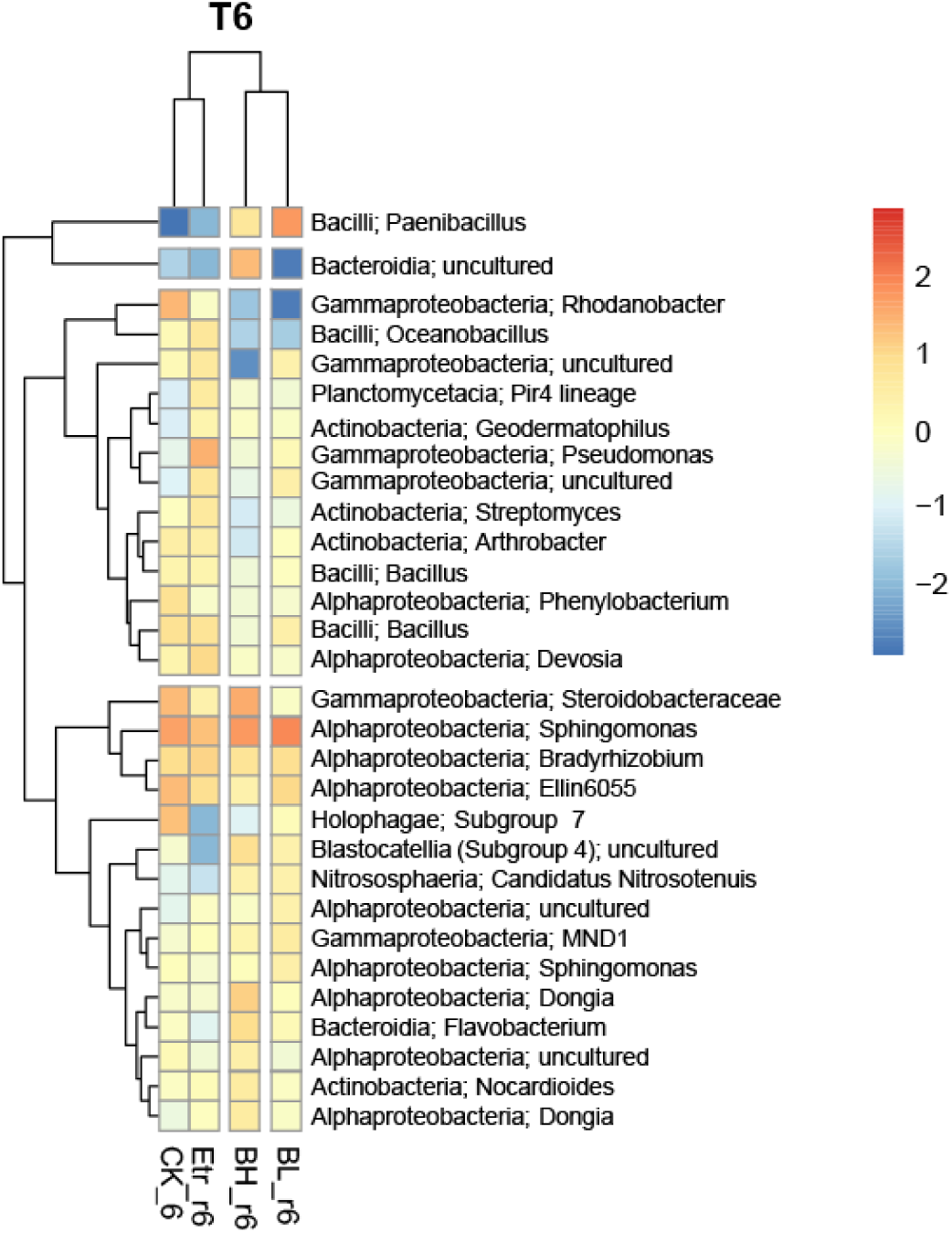
The relative abundance of the top 30 ASVs in the rhizome parts for four treatments in the latest sampling time. CK indicates ginger soil without any treatment, BL is low amounts of *B. velezensis*, BH is high amounts of *B. velezensis*, and Etr is the Etridiazole treatment.

At the first time point, *Bacillus* was found in four groups and was solely dominant in non-bacterial treatment groups; the second time point showed that the relative abundance of *Pseudomonas* had increased to become the dominant genus with *Bacillus*; at the third time point, *Bacillus* remained as a higher relative abundance genus; for the fourth time point, the relative abundances of *Sphingomonas* and *Paenibacillus* increased beyond that of *Bacillus. Sphingomonas* was found in all four groups while *Paenibacillus* was only observed in the bacterial treatment group. At the fifth time point, the bacterial treatment groups had greater relative abundances of *Bacillus* and *Sphingomonas* than did the control and Etr groups; the bacterial treatment groups were dominated by *Bacillus* and *Sphingomonas* at the final time point. In addition, *Pseudomonas* was solely dominant in the Etr group at every time point.

Ginger production was highest in the high concentration BV treatment group (BH) and lowest in the Etr group (Figure 5, Table S1). Although neither groups were significantly different from the control, the BH and Etr groups were significantly different to one another.

**Figure 5.**
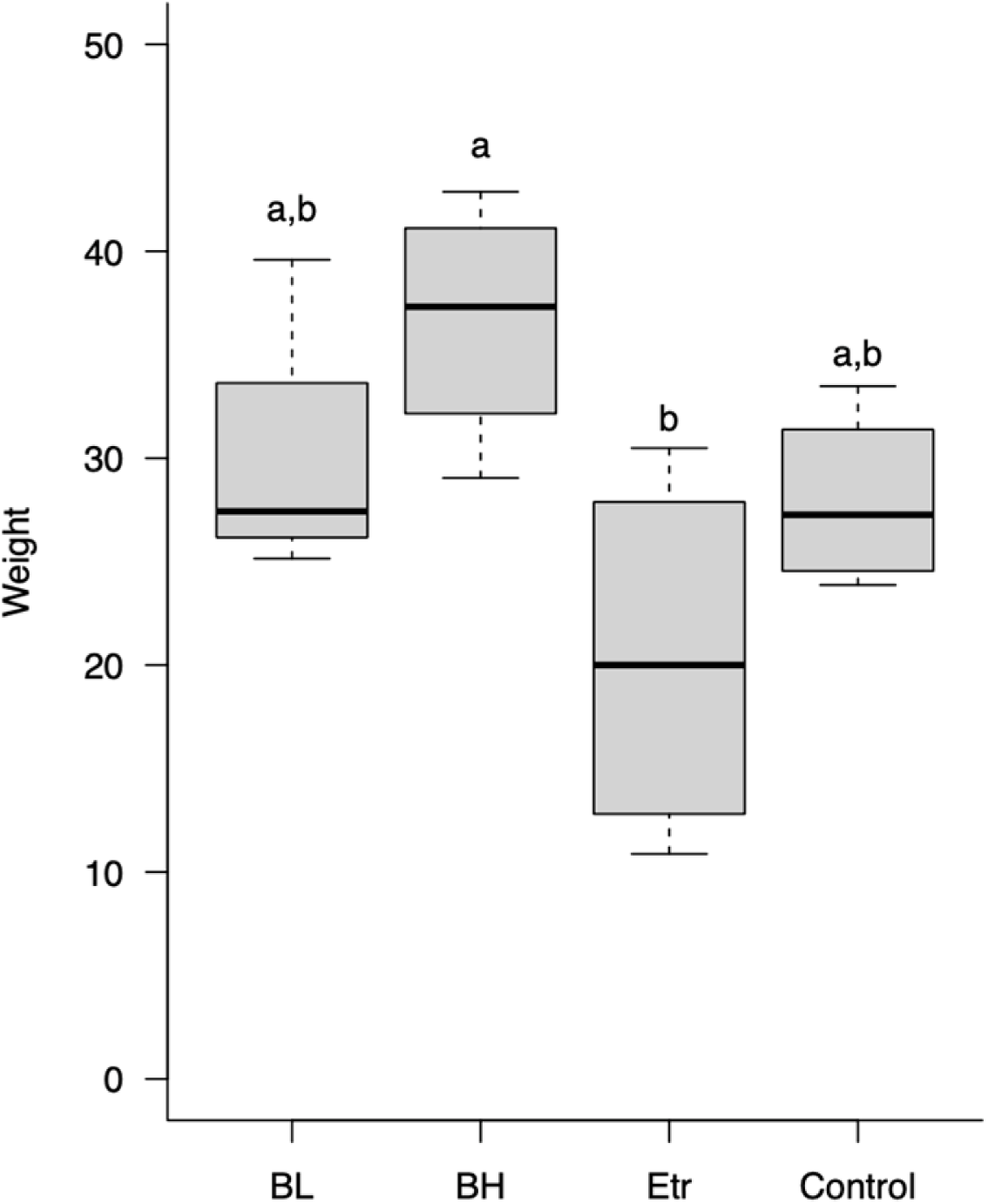
Production of gingers after different treatments. Control indicates ginger soil without any treatment, BL is low amounts of *B. velezensis*, BH is high amounts of *B. velezensis*, and Etr is the Etridiazole treatment.

To determine the relative abundance of *B. velezensis* (BV138) in the bacterial composition after adding it to the soil, we further compared all the sequences of *Bacillus* contained in the soil samples. The database contains 69 amplicon sequence variants (ASVs) that were *Bacillus*, and only three that were identified as *B. velezensis* (similarity of about 99%). These *B. velezensis* sequences were found at every time point among the BH samples and its average relative abundance was 0.24%. In the BL group, *B. velezensis* was mainly found at the third to the sixth time point of partial samples and its average relative abundance was merely 0.03%. There were no *B. velezensis* sequences in the control or Etr group. Based on the results, it indicates that *B. velezensis* (BV138) relative abundances in soil samples were positively correlated with its added amount, which is dose dependent.

## Discussion

Ginger is an important crop, but little research has been done on it compared to other agricultural plants. In the present study, we investigated the bacterial compositions in the rhizome and bulk regions of both healthy and diseased soils that previously grew ginger; we found that bacterial diversity was lower in diseased soil than in healthy soil, a phenomenon that was magnified in the rhizome region. Generally, the soil microbial community was influenced by plant growth because of the chemicals released by plant roots (17). Therefore, in the rhizome, microbial diversity decreased but microbial biomass increased in bulk soils (17). When the plant gets a disease, the microbial community dynamics may be caused by the plant. A review from (18) showed that plants can search for microbial assistance to resist pathogens.

In our study, the rhizome of the diseased soils showed an increase in the relative abundance of some bacteria, in addition to the pathogen *R. solani*. Members of *Flavobacterium* and *Chitinophagaceaea* had relatively higher abundances in diseased soil than healthy soil. Members of *Flavobacterium* are usually found in the rhizosphere with high abundance and have been thought to play a role in protecting plants from disease (19). A recent report indicated that *Chitinophaga* and *Flavobacterium* in endophytic bacterial communities may have the ability to suppress pathogens from soil (19). For example, sugar beet roots can attract *Chitinophaga* and *Flavobacterium* into the endosphere to suppress the fungal pathogen *R. solani* (20). Although our study did not investigate endophytic bacteria, the relative abundance of *Flavobacterium* did increase with that of *Ralstonia*, indicating that *Flavobacterium* may suppress *Ralstonia* in the root area of ginger. In addition, some species in *Stenotrophomonas* and *Sphingobacterium* were found to participate in inhibiting the growth and virulence of plant pathogens and have ability to rescue plants from stresses(21, 22). Both genera had higher relative abundances in the diseased soil than healthy soil in the present study. However, it is not clear if their abundance increased as a result of the plant host becoming infected.

Bacterial strains have been used as biofertilizers to ameliorate plant production, and chemicals have been used as pesticides and fungicides to maintain plant health. In this study, we found that applying the bacterial strain *B. velezensis* and fungicide Etridiazole can decrease bacterial compositions in soil, especially in the rhizome part. The influence is mainly constrained to the rhizome, which comes in contact with the ginger root, indicating that ginger may influence the bacterial composition during the treatments. As we discussed previously, plant roots may release some chemicals to adjust the soil microbial community (17).

However, in this study, we found that using Etridiazole results in the largest change in bacterial composition. Etridiazole (5-ethoxy-3(trichloromethyl)-1,2,4-thiadiazole) causes the hydrolysis of cell membrane phospholipids into free fatty acids and lysophosphatides, leading to the lysis of membranes in fungi. Therefore, it has been used as a fungicide. In addition to damaging fungi, etridiazole has side effects on other soil microorganisms because it reduces the nitrification rate of ammonium-oxidizing bacteria in soil, which may change the soil microbial community and influence the soil structure and function (23, 24).

In this study, *Pseudomonas* and *Bacillus* had higher relative abundances in the treatment with etridiazole. According to Shen et al. (2019), some rice endophytes, such as *B. aryabhattai, B. aryabhattai*, and *P. granadensis* can tolerate two or more fungicides, including etridiazole. In addition, they found that some strains may fix nitrogen, solubilize phosphorus, and produce indole acetic acid (IAA), which may promote plant growth and is believed to be a biofertilizer for rice (25). Here, we suggest that *Pseudomonas* and *Bacillus* show a similar tolerance to etridiazole when it is used on ginger. Hence, although we did not treat the ginger with Etridiazole and *B. velezensis* together, based on the dominant bacterial genera in the soil with Etridiazole, we suggest that *Pseudomonas* and *Bacillus* could be bacterial biofertilizers for ginger when used with the fungicide Etridiazole.

*Bacillus* species are PGPR that can form endospores that survive when dry into a dry powder for long shelf lives. Application of spore-forming *Bacillus* spp. does not have a lasting effect on the composition of the rhizosphere bacterial community (26). In this study, *B. velezensis* changed the bacterial compositions in soil, but *B. velezensis* can also increase the production of ginger in a dose-dependent manner.

Some mechanisms of *B. velezensis* have been studied in other plants. For example, *B. velezensis* FZB42 produces bioactive molecules that are active against microorganisms (27), including surfactin, iturin, and fengycin, all antifungal, lipopeptide compounds (28). Moreover, the antibacterial compounds difficidin and bacilysin are also produced by *B. velezensis* and are responsible for antagonistic activity against *Xanthomonas oryzae* pv. *oryzae* and *X. oryzae* pv. *oryzicola*, which cause rice diseases such as bacterial blight and bacterial leaf streak (29). *B. velezensis* also synthesizes plantazolicin, which kills parasitic nematodes (30). Furthermore, it was found that the biofilm formed by *B. velezensis* in plant rhizospheres can promote plant growth and secrete antimicrobial compounds to resist the invasion of infectious microbes (31).

Here, although the mechanisms are not yet clear, we found that the relative abundances of *B. velezensis* in samples were not high. We suggest that *B. velezensis* does not improve the ginger production by itself directly, but instead may influence ginger production indirectly by adjusting the bacterial community gradually. Bacteria belonging to the genus *Paenibacillus* have been isolated from diverse environments, especially from soils. Many of them ameliorate crop growth via biological nitrogen fixation, phosphate solubilization, and production of the phytohormone indole-3-acetic acid (IAA), and release of siderophores to increase iron acquisition (32). For example, *Paenibacillus jamilae* HS-26, which synthesizes hydrolytic enzymes and releases extracellular antifungal metabolites and volatile organic compounds—primarily, N, N-diethyl-1, 4-phenylenediamine—has highly antagonistic activity against several soilborne pathogens (33). Some *Paenibacillus* species are used in commercial biofertilizers, such as *P. macerans*, but their performance may be limited by soil pH, salinity, moisture content, and temperature (34).

In this study, the relative abundance of *Paenibacillus* and *Bacillus* increased in the treatments with *B. velezensis*. It is known that some *Bacillus* and *Paenibacillus* can elicit induced systemic resistance (ISR) similar to members of *Pseudomonas*, stimulating plant defense mechanisms against pathogens (35). Therefore, *Bacillus, Paenibacillus*, and *Pseudomonas* may have similar functions in soils, causing functional redundancy, which may explain why their relative abundances change so much.

## Conclusion

Although we did not study the function of *B. velezensis* in ginger disease, we demonstrated that *B. velezensis* benefits ginger reproduction. We also suggest that *B. velezensis* may influence the plant by adjusting the soil microbial composition. Regarding the function of Etridiazole in ginger reproduction, although treatment with Etridiazole did not significantly decrease the plant’s production, we suggest that Etridiazole may have some side effects on the gingers or beneficial bacteria in the soils that should be investigated further. Furthermore, whether using *B. velezensis* affects Etridiazole’s impact on reproduction is an interesting question. Our results provide clues for further investigations focusing on the interactions among the ginger, biocontrol agents, and fungicides.

## Funding

This work was financially supported by grants 110AS-1.3.2-ST-aM and 111AS-5.4.6-PI-P2 from the Council of Agriculture, Executive Yuan and Technology and Biodiversity Research Center, Academia Sinica in Taiwan.

## Acknowledgements

We greatly acknowledge Dr. Po-Yu Liu for his support with the bioinformatics. We also thank Noah Last of Third Draft Editing for his English language editing.w

---

**Figure S1.**
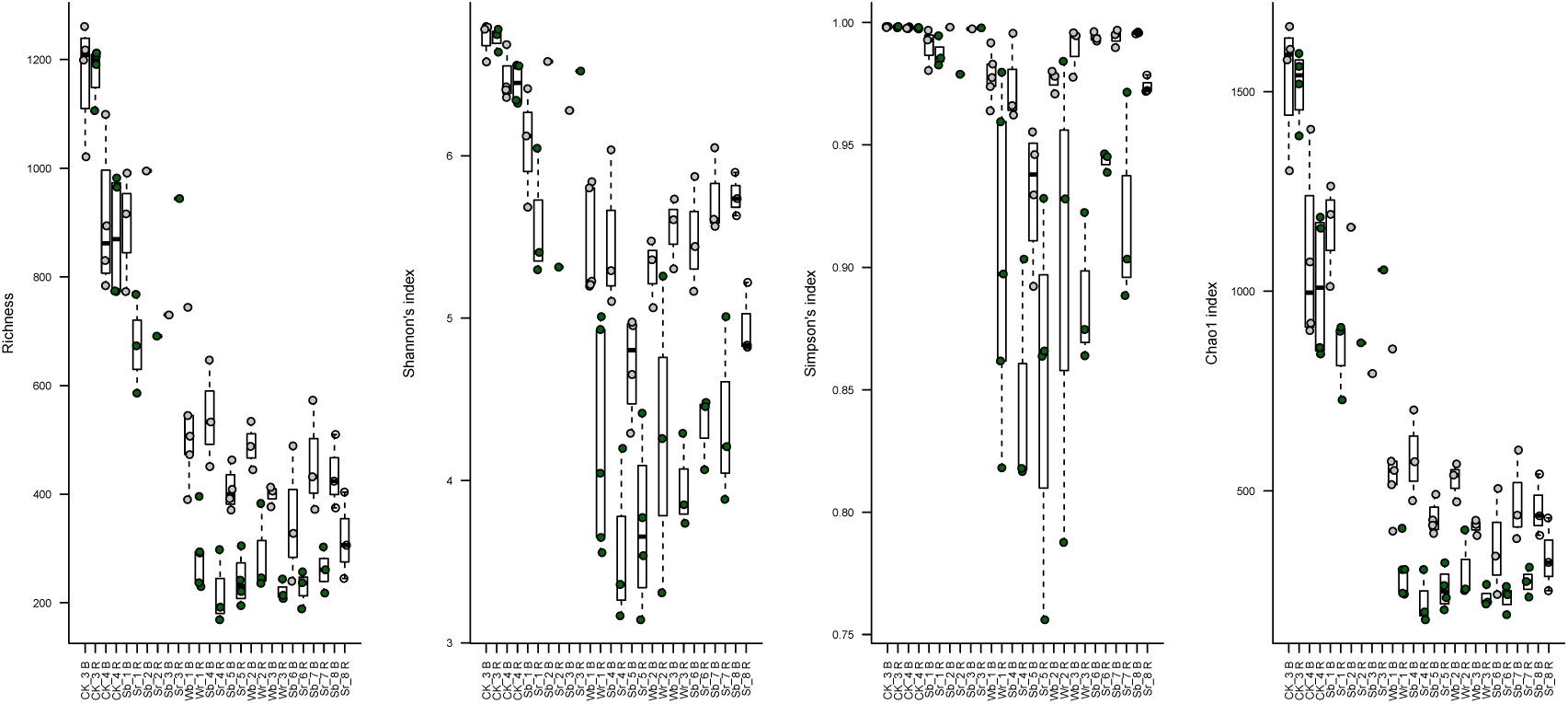
Alpha diversity indices of bacterial communities in each sample from bulk and rhizome parts of health and diseased soils. The grey color indicates the samples of bulk soil, and the green color indicates the samples of diseased samples. CK indicates control soil samples, Sr is soil with soft rot disease, and Wr is soil with wilt disease.

**Figure S2.**
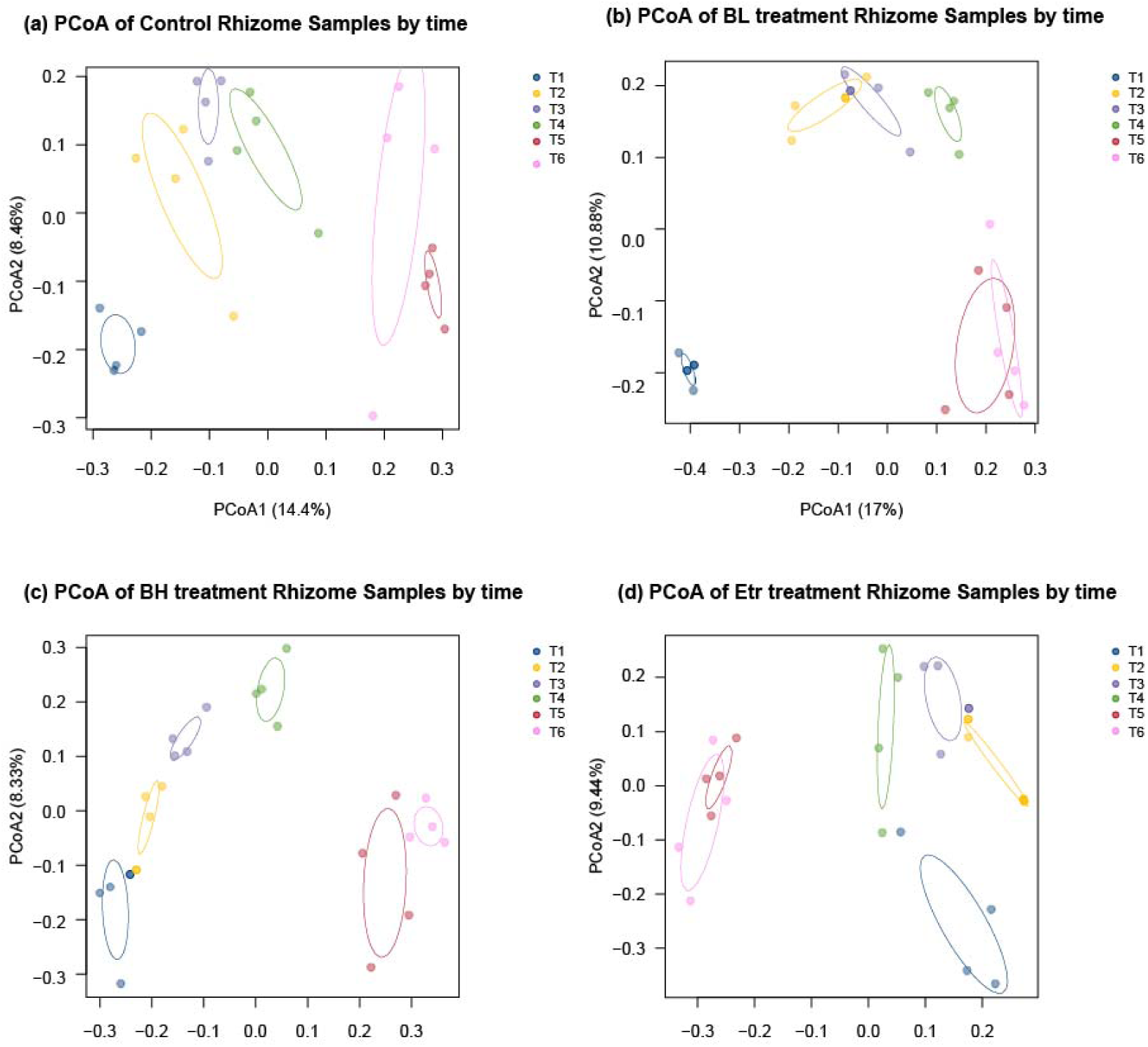
PCoA of bacterial communities from rhizomes in different treatments. Colors indicate the different time points.

**Figure S3.**
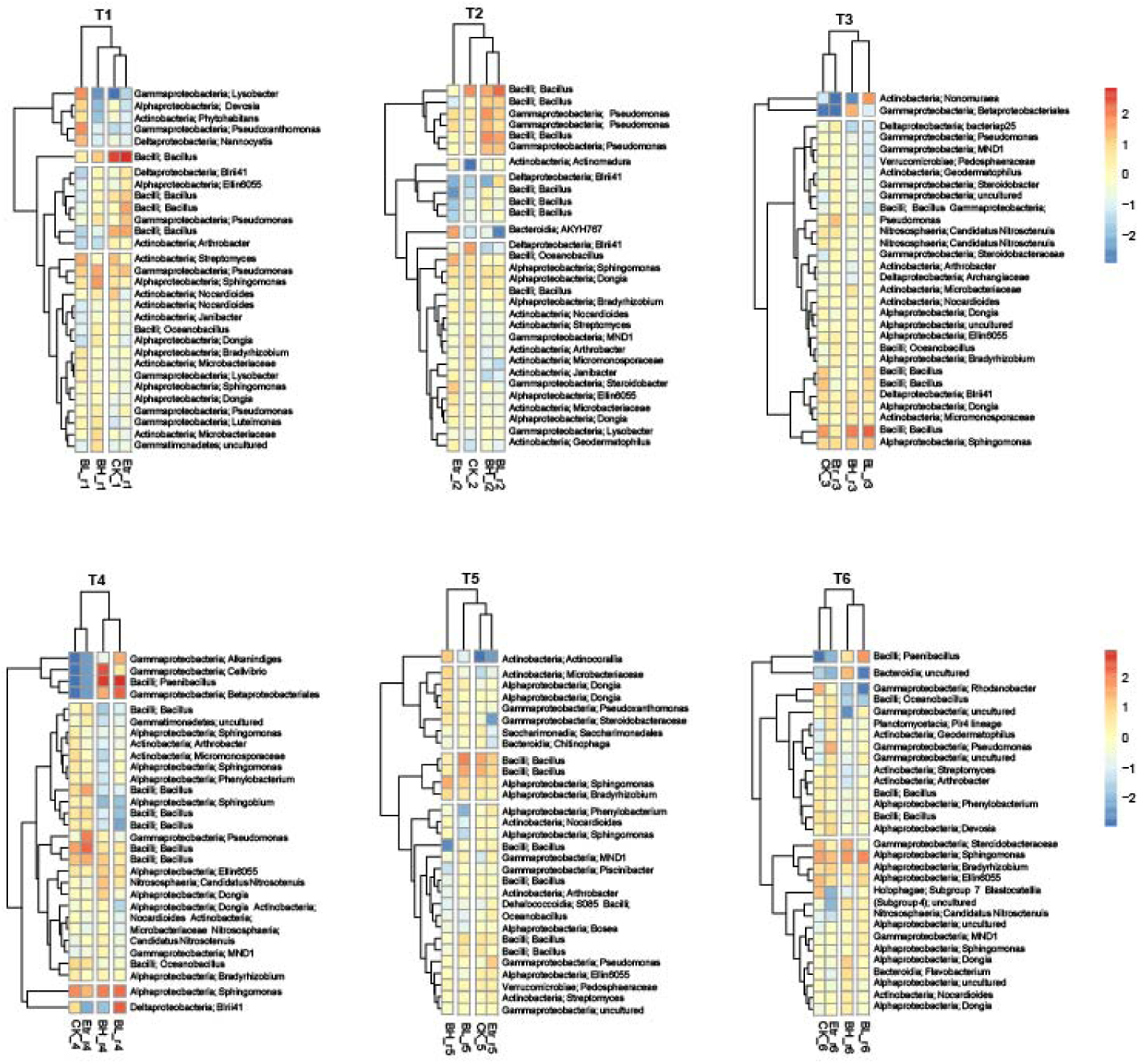
The relative abundance of the top 30 ASVs in rhizome parts for four treatments at each sampling time. CK indicates ginger soil without any treatment, BL is low amounts of *B. velezensis*, BH is high amounts of *B. velezensis*, and Etr is the Etridiazole treatment.

**Supplementary table 1.** Production information on the gingers in the study.

| Treatment | Duplicate1 | Duplicate2 | Duplicate3 | Duplicate4 | Average | SD | SE | Production (kg) | Ratio to Control |
| --- | --- | --- | --- | --- | --- | --- | --- | --- | --- |
| BL | 27.68 | 25.15 | 27.18 | 39.59 | 29.90 | 6.55 | 3.28 | 29.90 ± 3.28 ab | 1.07 |
| BH | 35.28 | 39.36 | 42.88 | 29.04 | 36.64 | 5.94 | 2.97 | 36.64 ± 5.94 a | 1.31 |
| Etr | 25.28 | 30.49 | 10.88 | 14.73 | 20.34 | 9.10 | 4.55 | 20.34 ± 4.55 b | 0.73 |
| Control | 29.30 | 25.23 | 23.87 | 33.48 | 27.97 | 4.34 | 2.17 | 27.97 ± 2.17 ab | 1.00 |

